# Biological rationales from language models enable leakage-resistant forecasts of target-indication success

**DOI:** 10.64898/2026.09.24.754137

**Authors:** Wanheng Zhang, Jie Xu, Zichen Zhang, Minghui Liu, Ryan Sun, Bissan Al-lazikani, Xiling Shen, Scott Kopetz, Lang Wu, Bingxin Zhao, Chong Wu

## Abstract

Identifying which target-indication (T-I) hypotheses can translate into clinical success remains a central challenge in drug discovery. We present PRIORITI (Prospective Rationale-Informed Outcome Reasoning for Integrated Target-indication Intelligence), a leakage-resistant framework that separates upstream biological evidence synthesis from outcome prediction. Given only a target gene and disease indication, a domain-instructed large language model (LLM) synthesizes drug-agnostic human genetics across prespecified channels. These evidence were embedded and combined with static entity context derived from gene summaries and disease definitions. The resulting representation is used to fit a supervised model trained on 6,784 temporally curated historical T-I outcomes, which is one of the largest cohorts assembled for this task. On a held-out historical test set of 1,696 T-I pairs, PRIORITI achieved a ROC-AUC of 0.883 and an expected calibration error (ECE) of 0.023, outperforming both a static target and indication context baseline and a human-curated genetic-evidence reference. On a strictly post-cutoff out-of-time benchmark of 765 T-I pairs, the largest such evaluation cohort reported to date, performance remained robust, with ROC-AUC 0.817, PR-AUC 0.357 and ECE 0.044, exceeding static target and indication context baseline and direct zero-shot LLM forecasting. A label-blinded LLM-based explainer further converted locked predictions into auditable prioritization rationales. Together, PRIORITI provides a calibrated, biology-first probability together with decision-oriented rationales for T-I prioritization before downstream translational investment.

## Introduction

Despite rapid advances in human genetics, molecular profiling and experimental biology, the conversion of biological insight into effective medicines remains slow, costly and failure-prone.^1–3^ The number of plausible therapeutic hypotheses has grown rapidly,^4^ but the capacity to evaluate them rigorously has not kept pace.^5^ The bottleneck is increasingly not hypothesis generation itself but prioritization, namely determining which of the many biologically plausible target-indication (T-I) relationships are supported strongly enough to justify substantial translational investment.

T-I prioritization is challenging because translational evidence is heterogeneous, incomplete and context-dependent.^6^ Human genetic evidence enriched in launched (i.e., success) T-I pairs^7,8^, but not necessarily the direction, tissue, timing or therapeutic tractability of modulation. Pathway biology may be coherent, yet too redundant, compensatory or pleiotropic to translate. Functional and animal studies may provide mechanistic support while imperfectly capturing human disease architecture.^9^ Conversely, an apparently modest signal in any single evidence channel may become compelling when integrated with convergent disease biology. Existing expert-curated resources and computational T-I algorithms have established the value of systematic evidence aggregation, including genetic support analyses^7,8^, target-disease association platforms^6,10,11^, and network-based indication-expansion methods^12,13^. These approaches are valuable, but they cannot fully solve this upstream prioritization problem at scale. Curated resources are labor-intensive to maintain and uneven across genes and indications, while many computational approaches compress complex biological arguments into annotations, scalar scores, graph features or latent factors. In parallel, models of drug approval or clinical-trial outcome often operate at the asset or trial level and may use molecule-, sponsor-, phase-, regulatory- or development-history features^14–16^ that arise after the original T-I decision. Such models address programme prognosis; they do not answer the earlier biological question of whether the underlying T-I hypothesis warrants pursuit.

Large language models (LLMs) offer a potential way to organize dispersed biomedical knowledge into explicit, pair-specific rationales.^17^ However, using them as direct forecaster creates a methodological problem. LLMs are probabilistic language models rather than calibrated predictive models.^18^ A single prompt asking whether a T-I pair will succeed conflates evidence retrieval, biological interpretation and probability estimation. The resulting judgement may be influenced by latent recall of known programmes, hidden training-data contamination, unverifiable evidence provenance ^19–21^ and poorly calibrated confidence^22^. These risks are particularly consequential in translational forecasting, where apparent accuracy may reflect recognition of prior development outcomes rather than prospective biological reasoning. A more principled strategy is therefore to constrain the LLM to construct an explicit, drug-agnostic biological argument from upstream evidence alone, and to separate this evidence-synthesis step from supervised learning of how recurrent evidence patterns relate to later clinical-development success.

Here, we present PRIORITI (Prospective Rationale-Informed Outcome Reasoning for Integrated Target-indication Intelligence), a leakage-resistant framework for forecasting T-I success from upstream biological evidence. To our knowledge, PRIORITI is the first framework to combine LLM-synthesized, drug-agnostic biological rationales, semantic embeddings of those rationales and supervised outcome learning at the T-I level. Given only a target gene and disease indication, a LLM (i.e., GPT-5) with domain instruction generates structured assessments and matched rationales across prespecified biological evidence channels, while excluding drug identities, trial outcomes, regulatory status, and other downstream development information. The resulting rationales are embedded together with target-function summaries and disease definitions, and a supervised model trained on temporally curated historical outcomes estimates the probability that at least one programme for the T–I pair will achieve clinical-development success. A separate label-blinded LLM explainer then converts each locked prediction into a decision-oriented biological rationale using only the evidence available at prediction time. We evaluated PRIORITI using a held-out historical test set, three entity-disjoint evaluations and a strictly post knowledge cutoff out-of-time benchmark. PRIORITI showed higher discrimination and better calibration than static target and indication context baselines, retained prioritization value under temporal separation and substantially exceeded direct zero-shot LLM forecasting. Most of the LLM-derived signal resided in the semantic structure of the synthesized rationales rather than in scalar evidence scores alone. These findings support an architecture in which LLMs synthesize inspectable biological evidence, whereas a supervised model maps recurring evidence patterns to calibrated probabilities of later clinical-development success, yielding a biology-first prior for early T-I prioritization.

## Results

### A leakage-resistant framework for auditable T-I success prediction

T-I prioritization is an upstream translational decision. At this stage, the relevant question is not whether a specific asset or trial will succeed, but whether modulation of a given target is biologically justified in a given disease context. We therefore formulated the task at the level of the T-I pair^8^. A pair was considered successful if at least one associated development programme reached launched, registered or pre-registration status; otherwise, the pair was classified as unsuccessful (**Methods**).

We developed PRIORITI to enforce a strict division between biological evidence synthesis and outcome modelling (**Fig. 1A**). For each T-I pair, we applied the domain-grounded instruction protocol using only the target gene and disease indication as inputs to the generative step (**Methods**). Under this protocol, GPT-5 synthesized drug-agnostic upstream human genetics and disease biology across four prespecified evidence channels, including human genetic causality, biological coherence, functional and animal support, and cross-source consistency. To minimize outcome circularity and downstream leakage, external tools were disabled and drug-, trial- and regulatory information were explicitly excluded from the instruction. The rationales were concatenated and embedded using a text embedding model, with Google Gemini Embedding 001 used in the primary analysis and several others in sensitivity analysis, to capture the semantic structure of the biological argument. This representation was intended to preserve information not reducible to scalar scores alone, including the relationship among evidence channels, the distinction between direct and indirect support, and how conflicting or incomplete evidence was reconciled. To encode baseline target and disease identity independently of the LLM reasoning step, we additionally embedded gene functional summaries and disease definitions (**Methods**). The final feature representation therefore combined scalar evidence scores, rationale embeddings, gene-summary embeddings and indication-definition embeddings. These features were then applied to a supervised predictor trained exclusively on historical outcomes (**Methods**). Unless otherwise stated, the primary predictor used throughout the study was a weighted AutoGluon^23^ ensemble combining LightGBM^24^, CatBoost^25^, RealMLP^26^, TabDPT^27^ and RealTabPFN-v2.5^28,29^. For each T-I pair, PRIORITI produced an estimated probability that the biological hypothesis would support at least one successful clinical-development programme.

**Fig. 1.**
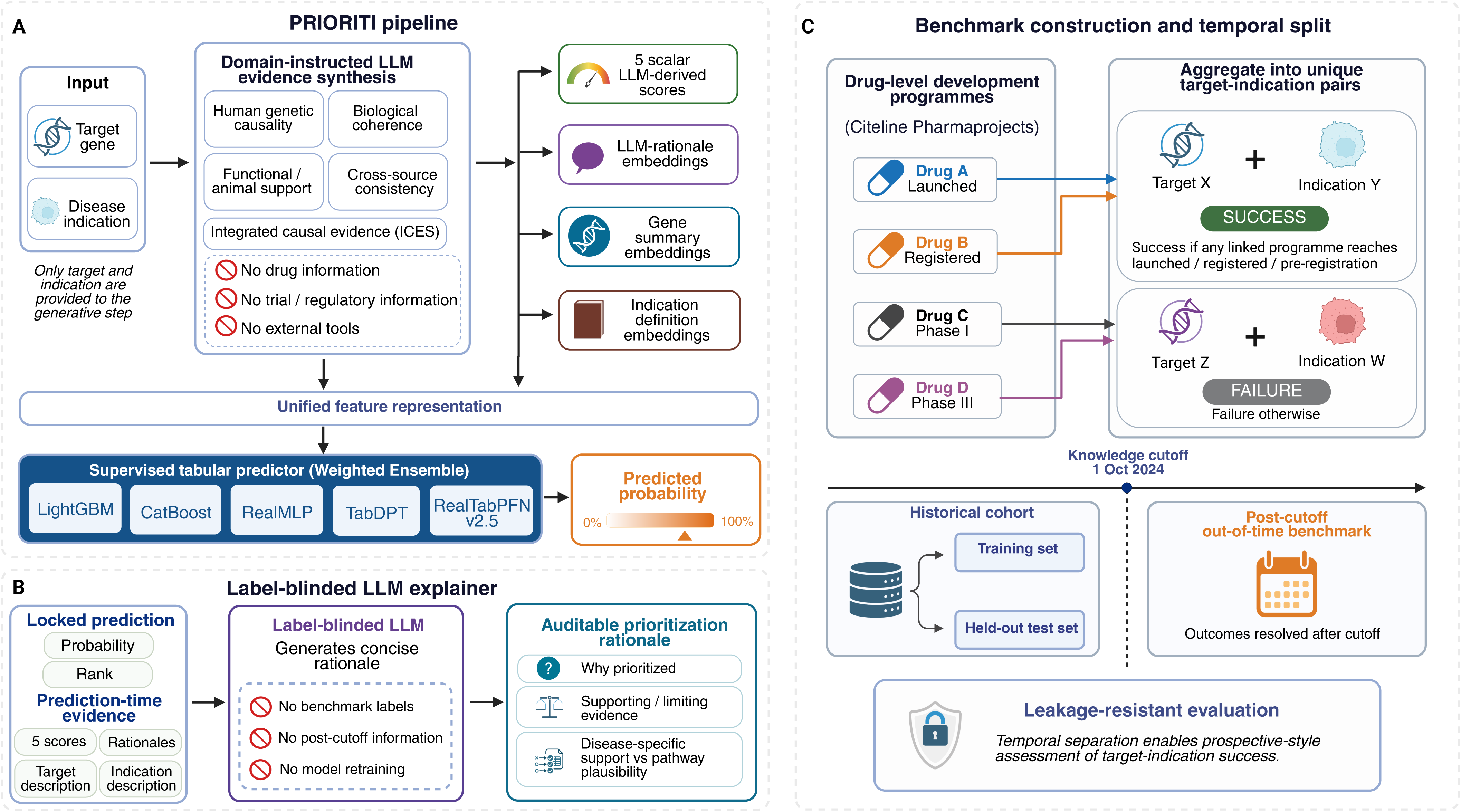
Study design of the PRIORITI framework for target-indication clinical development forecasting. **A,** Overview of the PRIORITI pipeline. A domain-instructed large language model receives only the target and indication as input and synthesizes evidence across prespecified biological channels, producing five scalar evidence scores and channel-specific rationales. These outputs are converted into a unified feature representation comprising scalar LLM-derived evidence scores, LLM rationale embeddings, gene summary embeddings and indication definition embeddings, which are then provided to a supervised tabular predictor to estimate the probability of T-I success. **B,** Label-blinded explanation of locked predictions. Each finalized prediction—its success probability and rank together with the prediction-time evidence (the five scalar scores, channel rationales, and target and indication descriptions)—is passed to a separate, label-blinded LLM that generates a concise rationale without access to benchmark labels or post-cutoff information and without any model retraining. The result is an auditable prioritization rationale that states why a pair was prioritized, summarizes the supporting and limiting evidence, and distinguishes disease-specific support from general pathway plausibility. **C,** Schematic of dataset construction at the target-indication (T-I) level. Drug-level development programmes from Citeline Pharmaprojects were aggregated into unique T-I pairs and assigned pair-level success or failure labels. The historical cohort was used for model development and split into training and held-out test sets, whereas T-I pairs with outcomes resolved after the October 1, 2024 knowledge cutoff were reserved as a post-cutoff out-of-time benchmark.

To ensure that narrative interpretation could not influence prediction, model training, operating-threshold selection and probability estimation were completed before the explainer was applied. The predicted probabilities and ranks were then frozen and passed to a separate, label-blinded LLM explainer (**Fig. 1B**). For each T-I pair, the explainer received only the information available at prediction time, including the predicted probability, cohort-relative rank, five evidence scores, matched channel-level summaries, and the target and indication descriptions (**Methods**). Its role was limited to translating the locked model output into a concise, decision-oriented rationale that identified the evidence supporting prioritization or deprioritization and distinguished disease-specific causal support from broader pathway plausibility.

We next constructed the T-I clinical-development outcome resource used for model development and evaluation. We curated Citeline Pharmaprojects records for monotherapy programmes that had advanced to at least Phase I and could be mapped to both a human gene target and an indication defined in the Medical Subject Headings (MeSH) ontology. Aggregating drug-level records and assigning binary pair-level outcomes yielded 23,463 unique T-I pairs and, to our knowledge, represents the largest clinical-development outcome dataset assembled (**Methods**; **Supplementary Table 1**). From this full resource, we derived two non-overlapping analytic cohorts using 1 October 2024, the knowledge cutoff of GPT-5, as the index date (**Fig. 1C**). The historical cohort was restricted to 8,480 T-I pairs whose outcomes had resolved before 1 October 2024 and that were also represented in Minikel et al.^8^ This restriction anchored model development to an established, mature benchmark and enabled direct comparison with prior genetic-support analyses. The historical cohort was randomly partitioned in an 80:20 ratio into a training set (N = 6,784) and a held-out test set (N = 1,696). The out-of-time cohort comprised T-I pairs absent from the Minikel et al.^8^ whose outcomes resolved only after the knowledge cutoff, yielding an independent post-cutoff benchmark of 765 newly resolved pairs (**Fig. 1C; Methods; Supplementary Tables 2 & 3**).

### LLM-synthesized rationales enable accurate and calibrated forecasting of T-I success

Because targets and indications recur across development programmes, T-I outcomes may be partly predictable from general target and disease characteristics alone. We therefore established a static entity-context baseline using only embeddings of existing gene functional summaries and standardized indication definitions, with all LLM-derived evidence excluded (**Methods**). This model used the same supervised-learning and evaluation pipeline as PRIORITI, providing a new pair-level benchmark for the signal contained in generic entity descriptions and isolating the additional value of LLM-synthesized evidence. Notably, a common-cohort comparison with published T-I and downstream clinical-trial forecasting models was not feasible: the former rely on separately curated target-disease evidence and outcome matrices, whereas the latter require extensive (and in some cases proprietary) molecular, trial-design, sponsor and longitudinal development data that were not available^14,16,30,31^ in harmonized form for the same pairs in our cohort.

On the historical held-out test set (N = 1,696; success prevalence, 13.5%), PRIORITI achieved strong discrimination, with a ROC-AUC of 0.883 (95% confidence interval (CI): 0.861–0.905) and a PR-AUC of 0.589 (95% CI: 0.526–0.656), compared with 0.863 (95% CI: 0.840–0.887) and 0.529 (95% CI: 0.462–0.596), respectively, for the static baseline (**Figs. 2A & 2B**). Results were stable across alternative hold-out partitions (**Supplementary Figure 1**). A reference model based on previously curated human genetic evidence^8^ provided little discrimination (ROC-AUC of 0.530; PR-AUC of 0.159).

**Fig. 2.**
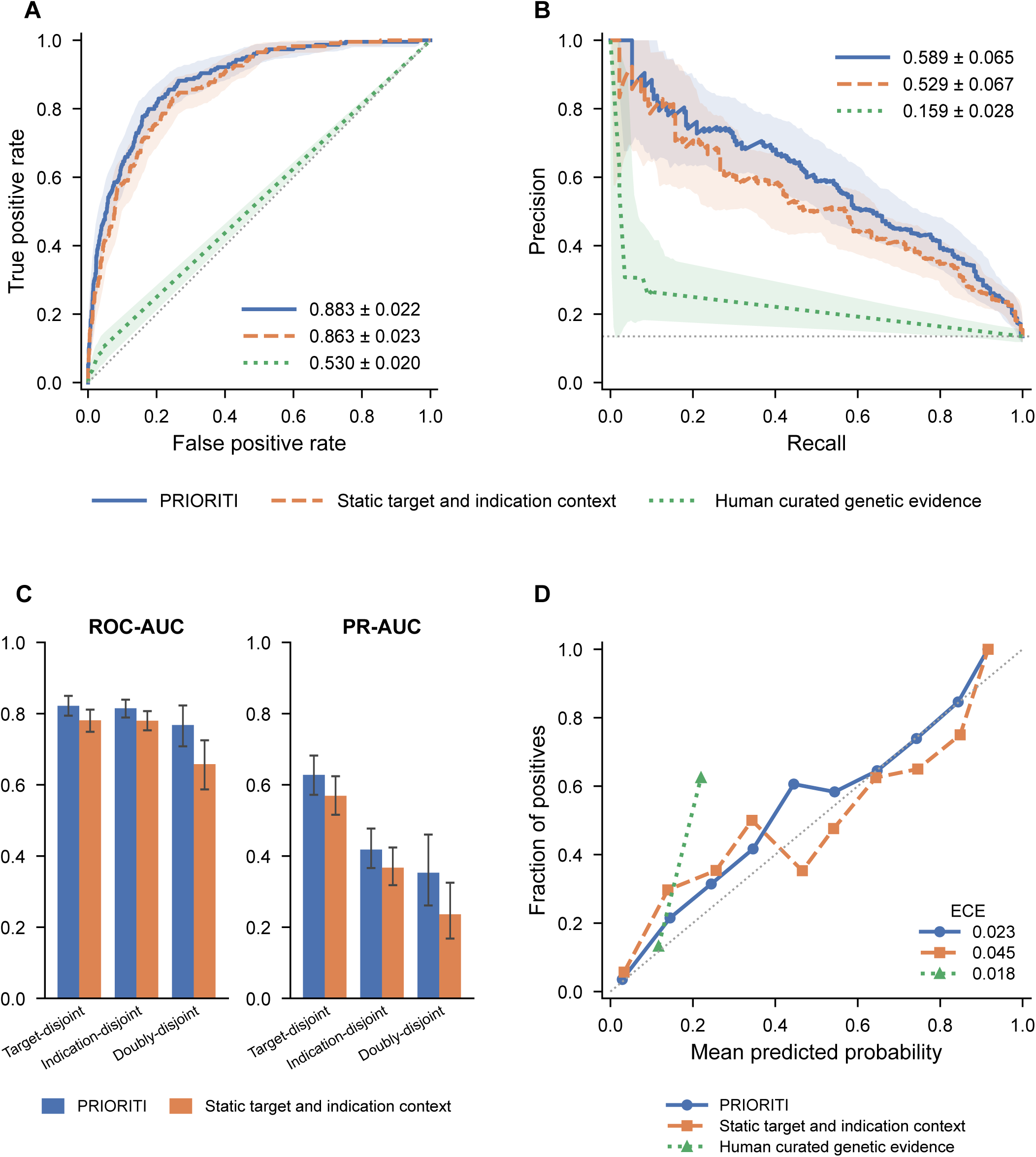
Historical held-out and entity-disjoint evaluation shows the discrimination and calibration of LLM-derived reasoning. **A,** ROC curves comparing the full PRIORITI model, a static target and indication context baseline (512-dimensional gene-summary and 512-dimensional indication-definition embeddings, with no LLM-derived evidence), and a reference model based on curated genetic evidence on the historical held-out test set. **B,** PR curves for the same three models on the historical held-out test set. **C,** Discrimination in three strict identity-disjoint evaluations (target-disjoint, indication-disjoint and doubly-disjoint), in which exact target and/or indication overlap between the training and test sets is removed, shown as ROC-AUC (left) and PR-AUC (right) for PRIORITI (blue) versus the static target and indication context baseline (orange); bars are point estimates and error bars are 95% confidence intervals. **D,** Calibration plot showing observed outcome frequency as a function of predicted probability for the three models on the historical held-out test set; ECE is indicated.

This random split, however, did not establish generalization to unseen biological entities. Although exact T-I pairs were held out, both the target and indication had each appeared in other training pairs for 94% of test pairs. We therefore retrained PRIORITI and the static baseline under target-disjoint, indication-disjoint and doubly-disjoint partitions (**Methods**; **Fig. 2C**). PRIORITI consistently outperformed the static baseline when targets were unseen (ROC-AUC, 0.822 versus 0.781; PR-AUC, 0.628 versus 0.569) and when indications were unseen (ROC-AUC, 0.815 versus 0.780; PR-AUC, 0.418 versus 0.367). In the doubly-disjoint evaluation, in which neither entity had appeared during training (N = 678; prevalence, 11.5%), PRIORITI achieved a ROC-AUC of 0.768 (95% CI, 0.708–0.823) and a PR-AUC of 0.353 (95% CI, 0.261–0.460), compared with 0.658 (95% CI, 0.587–0.725) and 0.236 (95% CI, 0.168–0.325) for the static baseline. Paired bootstrap confirmed improvement in both ROC-AUC (Δ = 0.109; 95% CI, 0.054–0.163) and PR-AUC (Δ = 0.117; 95% CI, 0.047–0.193). The larger gain after removing target and indication overlap indicates that the LLM-synthesized evidence representation captures transferable information about the T-I relationship beyond generic entity context.

Beyond its overall discrimination gain, PRIORITI improved both probability calibration and potentially support portfolio triage. On the held-out historical set, PRIORITI was better calibrated than the static target and indication baseline, with an expected calibration error (ECE) of 0.023 (95% CI: 0.016–0.040), compared with ECE of 0.045 for the baseline model (95% CI: 0.034-0.060; **Fig. 2D**). The model trained on human-curated genetic-evidence reference had a nominally lower ECE (0.018; 95% CI: 0.003–0.034), but this reflected a compressed probability range rather than useful risk stratification: no predicted probability exceeded 0.22, and discrimination remained close to random. The improved calibration of PRIORITI was accompanied by better prioritization of the highest-ranked T-I pairs. Among the top 3% of held-out predictions (probability cutoff = 0.659), 41 of 51 selected T-I pairs were true successes, corresponding to an observed success rate of 0.804 (95% CI: 0.647–0.883). In comparison, for the static target and indication baseline, the success rate among the top 3% of held-out predictions was 0.706 (95% CI: 0.588–0.882). Precision remained above 0.60 for all cutoffs from the top 1% to the top 10% (**Supplementary Figure 2**). Conversely, low predicted probabilities provided a strong deprioritization signal. Among the bottom 20% of predictions, the negative predictive value (NPV) was 0.997 (95% CI: 0.991–1.000), with 339 failures among 340 selected pairs (**Supplementary Figure 3**). These results support the use of PRIORITI not only to nominate high-priority hypotheses, but also to identify T-I pairs with weak prospects for translational success. For binary triage, we evaluated the prespecified operating threshold selected from cross-validated training predictions by maximizing F1-score. At this fixed threshold, PRIORITI operated in a high-specificity regime on the historical held-out test set, correctly classifying 1,381 of 1,467 unsuccessful pairs and 124 of 229 successful pairs. This corresponded to an accuracy of 0.887, balanced accuracy of 0.741, precision of 0.590, recall of 0.541, specificity of 0.941 and F1-score of 0.565. Because the acceptable trade-off between false-positive advancement and false-negative deprioritization depends on the intended translational use case (**Supplementary Figure 4**), our primary analyses rely on threshold-independent metrics (ROC-AUC, PR-AUC, ECE, and Precision@3%).

Ablation analyses localized this performance gain primarily to the semantic representation of the synthesized rationale (**Supplementary Table 4**). Among single component models, embeddings derived from the LLM-generated rationales for each T-I pair were the strongest individual predictor (ROC-AUC of 0.867; PR-AUC of 0.539), substantially outperforming gene embeddings alone (ROC-AUC of 0.775; PR-AUC of 0.353), indication embeddings alone (ROC-AUC of 0.668; PR-AUC of 0.280), and the scalar LLM evidence profile (ROC-AUC of 0.645; PR-AUC of 0.235). The full model nevertheless improved over rationale embeddings alone, indicating that target and disease context contributed complementary information once the synthesized argument was available. Grouped permutation analysis supported the same conclusion. LLM reasoning embeddings had the largest marginal importance (0.251), followed by gene summary embeddings (0.230) and indication summary embeddings (0.129), whereas the scalar LLM scores contributed comparatively little additional information (0.012) (**Supplementary Figure 5**).

### Out-of-time validation preserves calibrated prioritization under strict temporal separation

On this out-of-time cohort (N = 765; 89 successes, 676 failures; prevalence of 0.116), PRIORITI retained substantial discrimination, achieving a ROC-AUC of 0.817 (95% CI: 0.772-0.860) and PR-AUC of 0.357 (95% CI: 0.278-0.469; **Figs. 3A & 3B; Supplementary Table 5)**. The PR-AUC corresponded to a 3.1-fold enrichment over the cohort prevalence, indicating that the model continued to concentrate successful T-I pairs despite temporal separation. Predicted probabilities remained well calibrated, with an ECE of 0.044 (95% CI: 0.030-0.067; **Figure 3C**). PRIORITI also outperformed a static target and indication context baseline, which showed weaker discrimination (ROC-AUC of 0.787, 95% CI: 0.740-0.835; PR-AUC of 0.281, 95% CI: 0.223-0.373) and poorer calibration (ECE of 0.064, 95% CI: 0.047-0.088). In a paired bootstrap over the shared pairs, the full model’s advantage was statistically significant for PR-AUC (Δ = +0.076, 95% CI: +0.015 to +0.148) and for ROC-AUC (Δ = +0.029, −0.008 to +0.068).

**Fig. 3.**
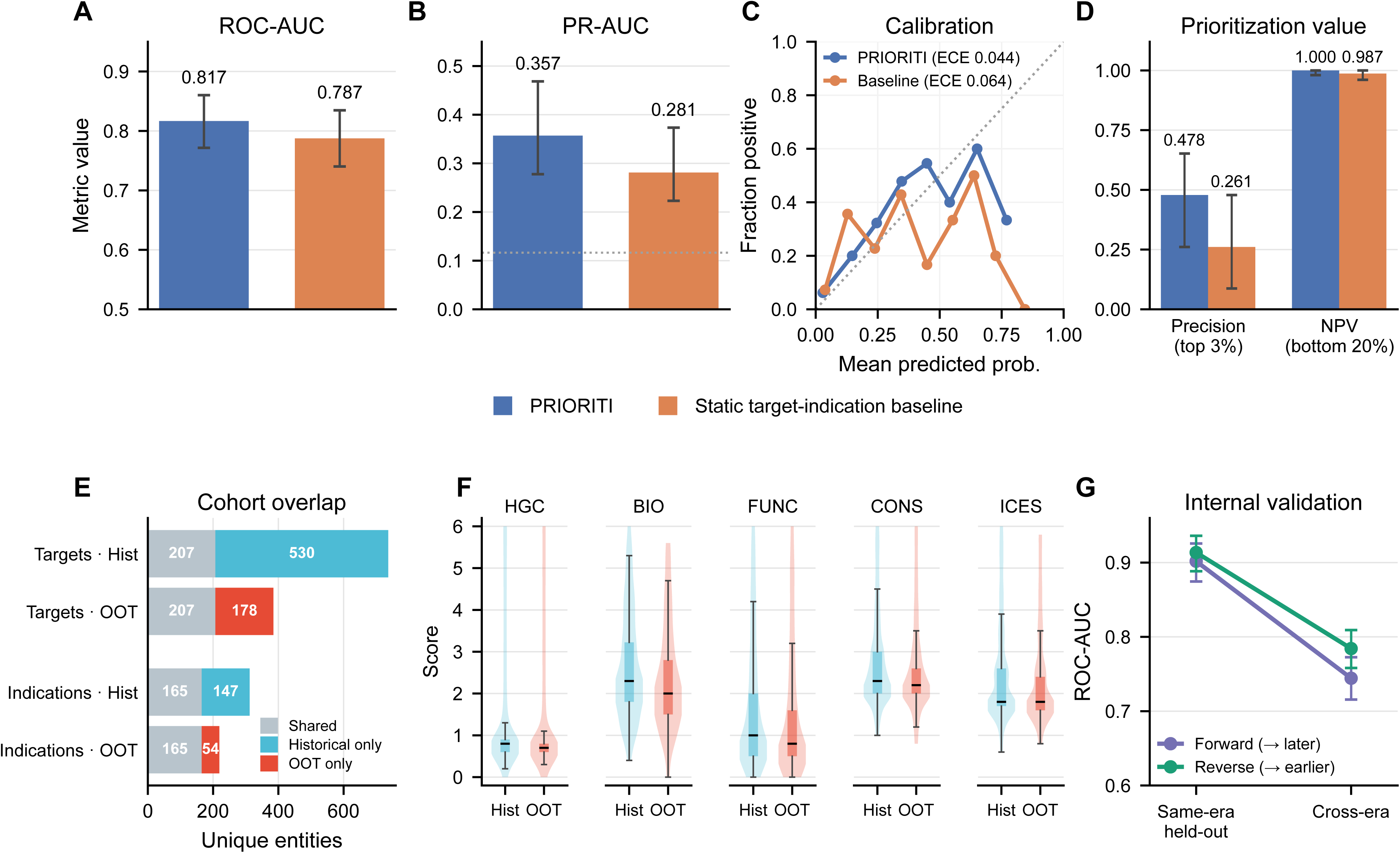
Out-of-time validation under temporal separation and analysis of cohort shift. A, ROC-AUC on the post-cutoff out-of-time (OOT) benchmark for PRIORITI and the static target and indication context baseline. **B,** PR-AUC for the same models; the dotted line marks the cohort success prevalence. **C,** Reliability (calibration) curves on the OOT benchmark, with ECE indicated. **D,** Prioritization value: observed success rate among the top 3% of predictions and NPV among the bottom 20%, for PRIORITI versus the baseline. **E,** Overlap in unique targets and indications between the historical and OOT cohorts, separated into shared and cohort-specific entities. **F,** Distribution of the five LLM-derived evidence scores (human genetic causality, HGC; biological coherence, BIO_COH; functional and animal support, FUNC_ANIMAL; cross-source consistency, CONSISTENCY; integrated causal evidence score, ICES) in the historical test and OOT cohorts. **G,** Internal temporal controls conducted entirely within the model’s knowledge horizon: a forward control (trained on earlier programmes, evaluated on a same-era held-out set and a later cross-era set) and a reverse control (trained on recent programmes, evaluated on a same-era held-out set and an earlier cross-era set), showing symmetric attenuation of ROC-AUC. Error bars, 95% bootstrap confidence intervals.

The ranked predictions retained practical prioritization value. Among the top 3% of out-of-time predictions, the observed success rate was 0.478, corresponding to a 4.1-fold enrichment over the cohort prevalence of 0.116, compared with 0.261 (2.2-fold) for the static target and indication context baseline (**Fig. 3D**). Conversely, the bottom 20% of predictions achieved a negative predictive value of 1.000 (95% CI 0.980–1.000), with all 153 selected pairs being failures, versus 0.987 (151 of 153; 95% CI 0.961–1.000) for the baseline. These results indicate that, even under temporal shift, high predicted probabilities enriched for later-resolved successes, while low probabilities provided a strong deprioritization signal.

Performance was lower than in the held-out historical test set, as expected for a genuinely forward-looking benchmark. The out-of-time cohort differed from the historical cohort in both entity composition and evidence strength. Target and indication overlap was only partial. The two cohorts shared 207 unique targets, whereas 530 were historical-only and 178 were out-of-time-only targets; similarly, 165 indications were shared, with 147 historical-only and 54 out-of-time-only indications (**Fig. 3E**). In addition, all LLM-derived evidence channels were modestly lower in the out-of-time cohort (**Fig. 3F**), as assessed by two-sided Mann-Whitney U tests that remained significant for every channel after Benjamini-Hochberg correction (all *P* < 10^-6^). It suggests that the post-cutoff benchmark contained, on average, less established upstream biological support. Together, these shifts indicate that the out-of-time benchmark represented a more difficult prediction distribution rather than a simple extension of the historical test set.

We next asked whether this attenuation reflected the loss of post-cutoff outcome information specifically, or the broader difficulty of transporting a model across development eras. To separate these explanations, we performed two internal controls conducted entirely within the model’s knowledge horizon (**Methods**). In a forward control, the model was trained on programmes resolved before 2022 and evaluated on a contemporaneous pre-2022 held-out set and on a forward set resolved between 2022 and the October 2024. ROC-AUC declined significantly from 0.902 (95% CI: 0.875–0.926) on the held-out set to 0.744 (95% CI: 0.716–0.773) on the forward set (Δ = 0.157, 95% CI: 0.120–0.194; **Fig. 3G**). In a complementary reverse control, the model was trained on recent programmes (resolved between 2015 and October 2024) and evaluated on earlier T-I pairs resolved between 2000 and 2014; performance again attenuated, from 0.914 (95% CI: 0.888–0.936) on a same-era held-out set to 0.784 (95% CI: 0.758–0.809) on the earlier set (Δ = 0.129, 95% CI: 0.093– 0.164). The comparable attenuation in both temporal directions argues against a simple explanation in which historical performance was driven primarily by latent access to known outcomes. Instead, it indicates that temporal distribution shift itself imposes a substantial generalization challenge for T-I forecasting.

### Direct zero-shot LLM forecasting underperforms supervised evidence mapping

Before comparing PRIORITI with direct LLM forecasting, we first assessed whether GPT-5 could recognize benchmark outcomes from prior knowledge. We gave GPT-5 only the target and indication names and asked whether each T-I pair represented a known success, a known failure, or an unknown outcome (**Methods**). Because the out-of-time benchmark comprises T-I pairs whose final outcomes were resolved only after the October 2024 knowledge cutoff, direct outcome recognition should be uncommon. Consistent with this expectation, GPT-5 labeled only 74 of 765 post-cutoff T-I pairs (9.7%) as already known. Manual adjudication against pre-cutoff Citeline records showed that 22 of these 74 putative recognition claims were not corroborated by pre-cutoff evidence; these were therefore classified as unsubstantiated and represented only 2.9% of the full benchmark cohort. The remaining 52 cases had pre-cutoff programme activity that could plausibly have supported partial label-relevant knowledge, and were therefore classified as “reasonable known.” However, even among these 52 cases, only 29 were directionally concordant with the final resolved outcome, whereas 23 were discordant **(Supplementary Table 6).** Thus, explicit outcome recognition by GPT-5 was infrequent, partly uncorroborated and imperfectly aligned with subsequent labels. This control supports the interpretation that the out-of-time benchmark primarily tests forecasting from structured biological evidence rather than simple recovery of known programme outcomes.

We next asked whether GPT-5 could directly forecast future programme outcomes in zero-shot sense. In the biology-only zero-shot setting, GPT-5 was asked to predict success or failure using only its internal biological and translational knowledge, without access to structured evidence summaries (**Methods**). Performance under this setting was limited, with a balanced accuracy of 0.550, recall of 0.146, and F1-score of 0.196 (**Fig. 4A; Supplementary Table 5**). We then evaluated an evidence-informed zero-shot setting in which GPT-5 was additionally provided with the same structured summaries and scalar evidence scores used to construct PRIORITI features (**Methods**). This improved direct forecasting modestly, with balanced accuracy increasing to 0.576, recall to 0.191, and F1-score to 0.259 (**Fig. 4A**), but performance remained clearly below that of PRIORITI. Directly prompted probabilities were also less well calibrated, with ECE of 0.127 for biology-only zero-shot and ECE of 0.103 for evidence-informed zero-shot, compared with 0.044 for PRIORITI on the same benchmark. These results clarify the role of the LLM in the framework. Providing structured evidence improved direct LLM judgment, indicating that the synthesized evidence was informative. However, the larger gain emerged only when these evidence patterns were embedded and mapped to historical T-I outcomes by a supervised predictor. In this setting, the LLM contributes more effectively as an evidence synthesizer than as a standalone forecaster.

**Fig. 4.**
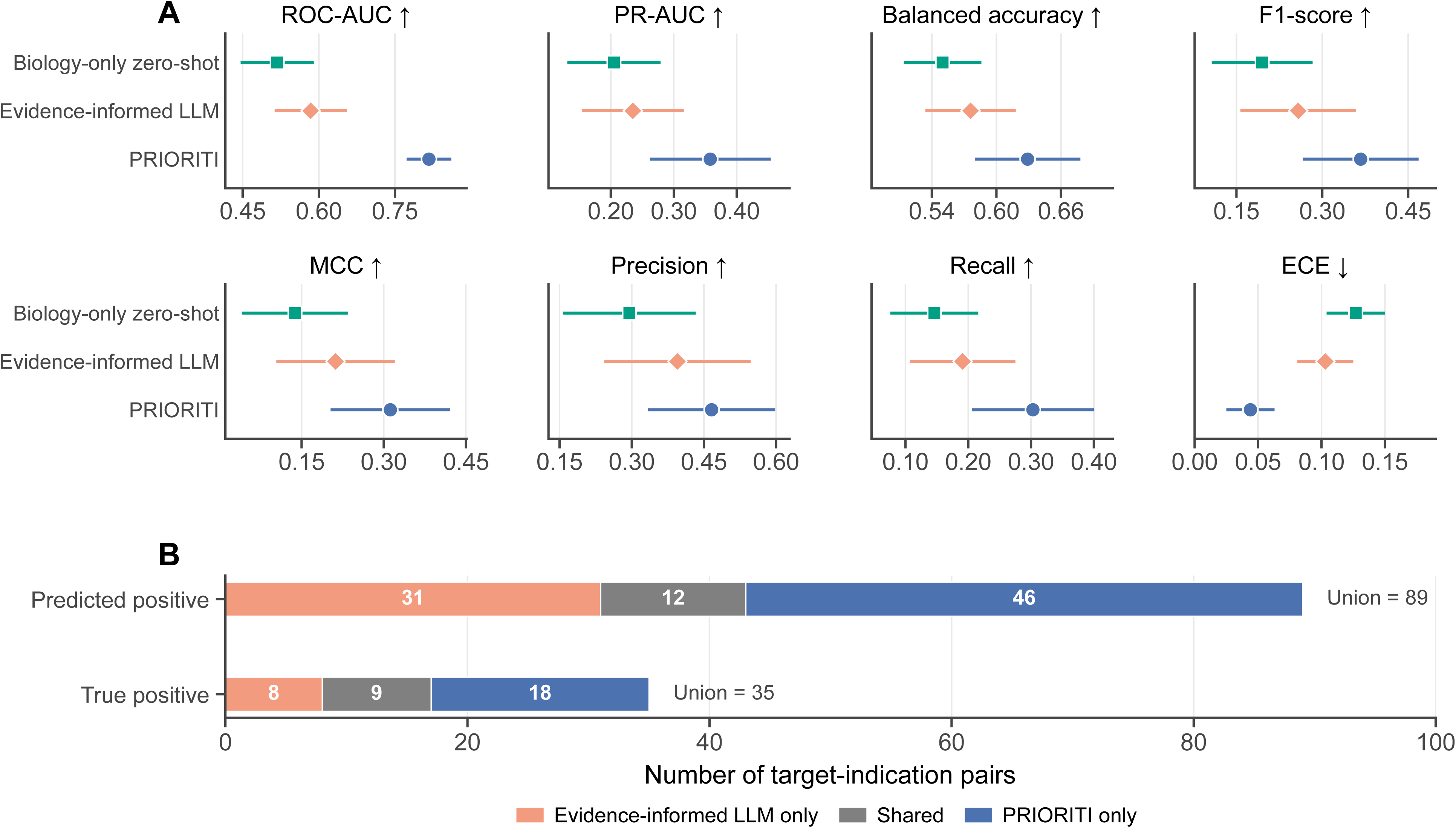
Direct zero-shot LLM forecasting underperforms PRIORITI but captures partially non-overlapping signal. **A,** Performance comparison of biology-only zero-shot LLM forecasting, evidence-informed zero-shot LLM forecasting, and PRIORITI on the post-cutoff benchmark across discrimination, threshold-based and calibration metrics. **B,** Overlap between evidence-informed zero-shot LLM predictions and PRIORITI predictions, shown for all predicted positives and for true positives only. Bars indicate model-specific and shared T-I pairs.

Despite weaker overall performance, the evidence-informed zero-shot forecast showed only limited overlap with PRIORITI, suggesting partial complementarity. PRIORITI classified 58 T-I pairs as success, whereas the evidence-informed LLM classified 43 as success, with only 12 pairs overlapping (**Fig. 4B**). Among true positives, 9 successful pairs were identified by both approaches, 18 were unique to PRIORITI, and 8 were unique to the evidence-informed LLM (**Fig. 4B**). This limited overlap suggests that direct LLM judgment may capture a small non-redundant component of signal. However, the larger number of unique true positives recovered by PRIORITI, together with its stronger overall performance and calibration, indicates that supervised evidence mapping remained the more reliable forecasting strategy.

Finally, we asked whether PRIORITI’s advantage persisted after removing all pairs that GPT-5 had considered potentially known. On the subset labeled unknown in the knowledge-recognition control (N = 691), PRIORITI retained superior predictive performance, achieving a balanced accuracy of 0.632, F1-score of 0.367, precision of 0.458, and recall of 0.306, whereas biology-only zero-shot forecasting remained weak (balanced accuracy = 0.532, precision = 0.259, recall = 0.097, F1-score = 0.141) and evidence-informed zero-shot forecasting, although improved, still substantially underperformed the trained predictor (balanced accuracy = 0.559, precision = 0.435, recall = 0.139, F1-score = 0.211) (**Supplementary Table 7**). Calibration also remained superior for PRIORITI on this subset (ECE = 0.038 versus 0.129 and 0.104, respectively).

### Case-level rationales explain PRIORITI predictions and define their biological boundary conditions

To examine whether PRIORITI forecasts could support practical T-I prioritization, we built a constrained, label blinded LLM-based explainer to all 765 out-of-time predictions after probabilities and ranks had been locked (**Methods; Supplementary Table 13**). For each T-I pair, the explainer received the locked probability and cohort-relative rank, the five LLM-derived evidence scores, their corresponding channel summaries, and descriptions of the target and indication. Benchmark labels, post-cutoff outcomes, drug identities, trial histories and regulatory information were withheld. The resulting rationales could not modify the predictions, and benchmark labels were introduced only after rationale generation to identify concordant and discordant cases. Because the explainer did not receive local attribution information from the fitted model, these outputs should be interpreted as evidence-grounded contextualization of the forecasts rather than causal explanations of how the model produced them.

Among concordant positive cases, the clearest recurring pattern was a disease-proximal mechanism connecting target function to pathophysiology. For example, for SLC5A2-Diabetes Mellitus (PRIORITI predicted probability of success of 0.638), the explainer linked renal glucose reabsorption as a direct mechanism linked to diabetic physiology. For PLG-Blood Coagulation Disorders (predicted probability of success of 0.776), the explainer connected plasminogen-mediated fibrinolysis to clot resolution and coagulation balance. In both cases, the translational chain was comparatively short in which the molecular function of the target mapped directly onto a disease-defining physiological process and an expected consequence of therapeutic modulation.

Other concordant cases illustrated how convergent evidence could support prioritization even when no single evidence channel was uniformly strong. IL12B–juvenile arthritis received a positive forecast (predicted probability of success of 0.647) despite limited direct human genetic support and only moderate integrated causal evidence. The explainer placed this limitation alongside convergent support from IL-12/IL-23 biology, disease-relevant immune cell context and functional evidence from inflammatory arthritis models. By contrast, MAPKAPK2-Inflammation received a negative forecast with a low predicted probability of success (0.013) despite very strong biological coherence and functional support from the model inputs. Here, the explainer emphasized that broad involvement in inflammatory signalling was not accompanied by comparably strong human genetic or indication-specific translational evidence. The contrast between these cases shows how similar strengths in pathway or functional evidence can lead to different forecasts when the broader configuration of evidence differs.

TNF-Heart Failure provided a more clinically established example of the distinction between pathway involvement and therapeutic causality. Despite strong biological coherence and functional and animal model support, the pair received negative forecast (predicted probability of success of 0.165). The explainer emphasized weak human genetic support and the gap between inflammatory pathway plausibility and disease-specific therapeutic translation in chronic heart failure. This interpretation is consistent with prior anti-TNF experience in heart failure, including the ATTACH trial^32^, in which high-dose infliximab increased the combined risk of death or heart-failure hospitalization, and RENEWAL^33^, in which etanercept was stopped early for lack of benefit. These trial results were not provided to the explainer. The case therefore illustrates why biological participation in a disease process does not necessarily imply that therapeutic modulation of the pathway will be beneficial.

Discordant cases highlighted that clinical success can arise through mechanisms that are not adequately represented by a single T-I pair. GLP1R-Sleep Wake Disorders and GIPR-Sleep Wake Disorders were both false negatives, receiving low predicted probabilities of 0.038 and 0.065, respectively, despite the subsequent approval of tirzepatide for obstructive sleep apnea in adults with obesity^34^. The explainer attributed the low model priority to limited evidence for a direct role of GLP1R or GIPR in sleep-wake disorder biology. This assessment is biologically coherent at the single-target level. tirzepatide is a dual GLP-1/GIP receptor agonist, and its benefit in obstructive sleep apnoea is likely mediated substantially through weight loss and broader cardiometabolic effects rather than through direct modulation of sleep–wake biology^35,36^.

To save space, we have relegated all other cases into the **Supplementary Table 13.** Collectively, these cases delineate the scope of PRIORITI. Its forecasts were most readily interpretable when target modulation could be connected to disease through a direct, indication-specific biological chain or through convergence across complementary evidence channels. The framework was less complete when clinical benefit depended on intermediate phenotypes, multi-target pharmacology, drug-specific properties, patient selection or indication reframing. Case-level explainer can therefore support biological review and hypothesis triage, but they should not substitute for asset-level assessment or expert translational judgment.

### Robustness analyses show that the forecasting signal is stable across learners, text embedding models and disease areas

We first assessed whether PRIORITI performance depended on a particular tabular learner. Across five distinct model families used within the AutoGluon framework, performance varied only modestly, with ROC-AUCs ranging from 0.858 to 0.883 and PR-AUCs from 0.507 to 0.589 (**Figure 5A**). The ensemble performed best (ROC-AUC of 0.883, 95% CI: 0.860 to 0.905; PR-AUC of 0.589, 95% CI: 0.525 to 0.655), but TabDPT, RealMLP, RealTabPFN-v2.5, Catboost and LightGBM all retained similar discrimination. This narrow performance range indicates that the forecasting signal was not an artefact of a single model class, but was recoverable across distinct supervised learning architectures when supplied with the PRIORITI representation.

**Fig. 5.**
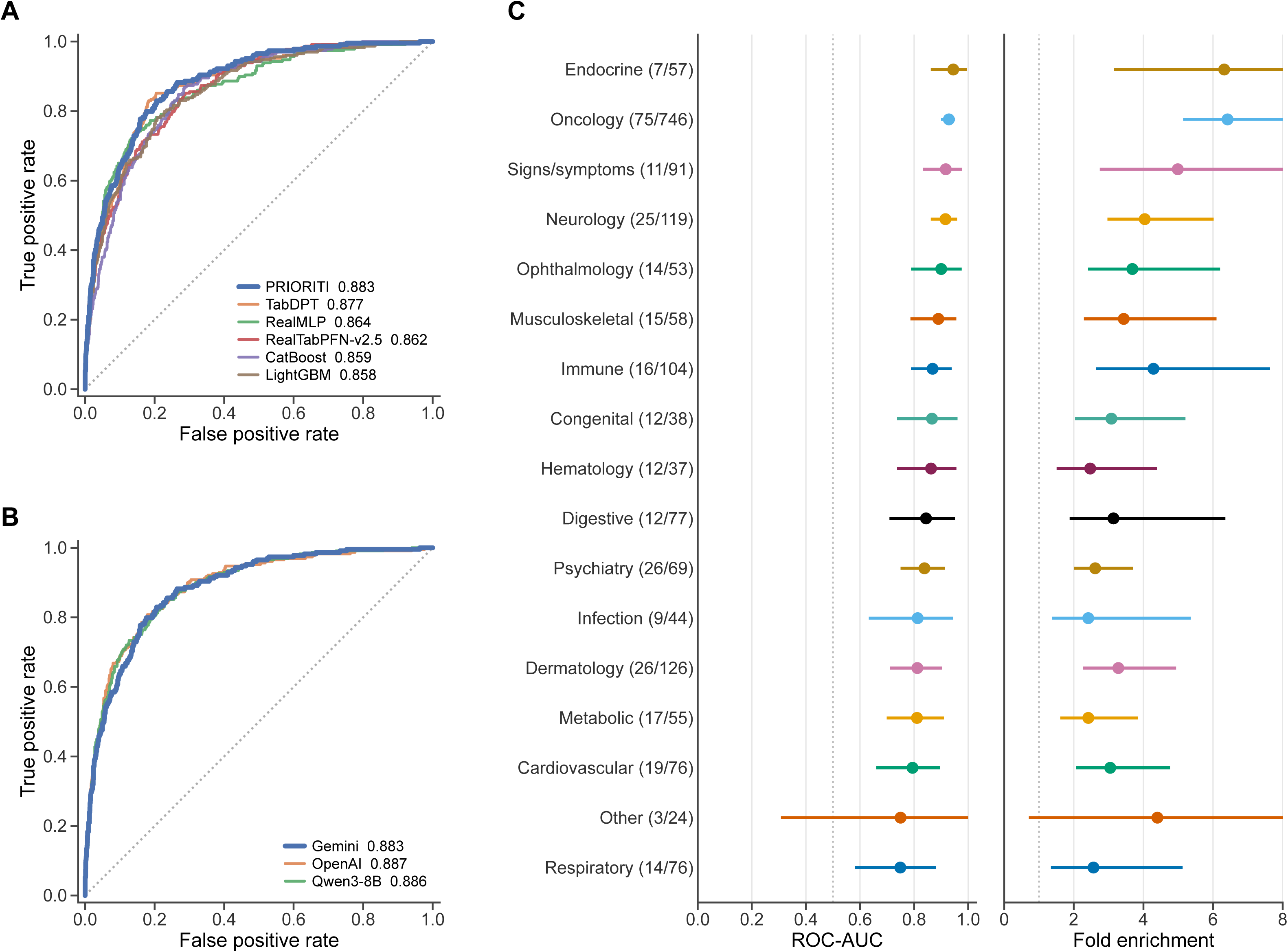
Forecasting performance is stable across learners, therapeutic areas and text-embedding models. **A**, ROC curves on the historical held-out test set for PRIORITI (the weighted AutoGluon ensemble) and the individual tabular learner families (TabDPT, RealMLP, RealTabPFN-v2.5, CatBoost and LightGBM); ROC-AUC is shown in the legend. **B**, Robustness to the choice of text-embedding model: ROC curves on the historical held-out test set for the full PRIORITI model built with Google gemini-embedding-001, OpenAI text-embedding-3-large, or Qwen3-Embedding-8B embeddings, with the feature set and embedding dimensionality held fixed (ROC-AUC shown in the legend). **C**, Performance stratified by therapeutic area, shown as ROC-AUC and fold enrichment over the within-area baseline prevalence for each disease category; numbers in parentheses indicate successes over failures in each stratum, and error bars are 95% bootstrap confidence intervals.

We then tested whether PRIORITI depended on the particular text-embedding backbone used to encode rationales, gene summaries and indication definitions. Holding the downstream architecture, feature set and embedding dimensionality fixed, we regenerated all text representations using Google gemini-embedding-001 (used in the main analysis), OpenAI text-embedding-3-large, and the open-weight Qwen3-Embedding-8B (**Methods**). Discrimination was essentially unchanged across backbones, with ROC-AUCs of 0.883 (95% CI 0.861–0.905), 0.887 (95% CI 0.863–0.911) and 0.886 (95% CI 0.863–0.908), and PR-AUCs of 0.589 (95% CI 0.526–0.656), 0.612 (95% CI 0.547–0.677) and 0.598 (95% CI 0.531–0.665) for Gemini, OpenAI and Qwen, respectively (**Figure 5B**). Calibration was also stable, with ECE of 0.023, 0.024 and 0.031, respectively. These results indicate that PRIORITI’s predictive signal was not dependent on a single embedding provider or representation model.

We then examined whether performance was concentrated in a small number of therapeutic areas or generalized across disease contexts. Stratification by therapeutic category showed consistently strong discrimination across contexts, with ROC-AUCs ranging from 0.749 to 0.945 (**Figure 5C**). Because PR-AUC depends strongly on outcome prevalence, we summarized precision-recall performance as fold enrichment over the within-area success prevalence (PR-AUC / prevalence). By this prevalence-adjusted measure, all therapy areas showed meaningful fold enrichment, with point estimates ranging from 2.42-fold to 6.42-fold over baseline. The strongest signals were observed in oncology (ROC-AUC of 0.929, 95% CI: 0.900–0.951; enrichment 6.42×, 95% CI: 5.14–8.09×) and endocrine disease (ROC-AUC of 0.945, 95% CI: 0.862–0.996; enrichment 6.32×, 95% CI: 3.15–16.0×). Respiratory disease showed the lowest point estimate, but still retained separation from baseline (ROC-AUC of 0.749, 95% CI 0.581–0.881; enrichment 2.57×, 95% CI 1.34–5.12×). Thus, although performance varied across disease contexts, the forecasting signal was not restricted to a single dominant therapeutic area.

## Discussion

In this study, we show that dispersed upstream human genetics and disease biology can be converted into a learnable, leakage-resistant representation for T-I prioritization. PRIORITI is built around a simple but important separation: the LLM synthesizes drug-agnostic biological evidence into explicit rationales, whereas the supervised model learns how recurrent patterns in those rationales relate to later T-I outcomes. PRIORITI improved on static target and indication context in the conventional held-out evaluation. Its advantage widened when targets and indications were both unseen during training and persisted in the out-of-time cohort, particularly for enrichment among the highest ranked hypotheses. Direct zero-shot LLM forecasting was less accurate and less calibrated, even when supplied with the same evidence summaries.

This separation design changes the role of the LLM in translational forecasting. Directly asking an LLM whether a T-I pair will succeed combines evidence retrieval, biological interpretation, latent memory, and probability estimation within a single opaque judgment, making it difficult to distinguish prospective reasoning from recall of known development history, hidden data contamination or poorly calibrated confidence. PRIORITI instead uses the LLM to construct an explicit intermediate representation of the biological evidence. This design does not guarantee that every generated claim is correct, but it makes the evidence structure available for inspection, comparison and empirical testing. The supervised predictor then estimates how recurrent patterns in that representation are associated with observed development outcomes, rather than relying on the LLM’s self-assessed confidence. Separating the post hoc interpretation layer (i.e., LLM explainer) from the locked prediction further prevents a plausible narrative from feeding back into and potentially altering the forecast.

The entity-disjoint evaluations help clarify what information this representation contributes. Static embeddings of gene summaries and disease definitions already captured substantial predictive signal, reflecting target identity, broad biological function, and disease taxonomy. Indeed, the relatively modest improvement in the conventional held-out set indicates that much of the apparent predictability in a random split can arise from familiar entities as 94% of held-out pairs shared both their target and indication with at least one training pair. The doubly-disjoint evaluation was therefore more diagnostic. When neither the target nor the indication had been observed during training, the advantage of PRIORITI over static context increased substantially. The out-of-time benchmark provides a complementary result. Even under temporal shift, PRIORITI improved discrimination, calibration and top-ranked enrichment over static context baseline. These findings suggest that the synthesized biological argument carries transferable information about T-I plausibility, not merely a memorized representation of known targets or indications.

PRIORITI also differs from many previous drug-development forecasting models in the decision context it is intended to support. Previous predictive models have often incorporated asset-, trial-, sponsor-, regulatory- or development-history features, which can improve retrospective discrimination, but may partially reflect information generated after the original prioritization decision.^14,37^ Such features are useful for forecasting the fate of an existing programme, but they address a different downstream question that assesses whether a particular programme is likely to succeed given what is already known about its execution. By contrast, PRIORITI was designed to address an earlier question that whether the biological relationship between a target and an indication is sufficiently supported to justify further translational investment before substantial asset-specific evidence accumulates. By excluding drug-, trial- and regulatory-level information from the evidence-synthesis step, the framework estimates a biology-first prior over T-I opportunities rather than a forecast of programme execution. This framing makes the method most relevant to target prioritization^6^, indication expansion^38^ and portfolio triage^39^.

The case-level analyses illustrate both the value and the limitations of PRIORITI forecast. Concordant positive examples were characterized by either a short mechanistic chain from target function to disease pathophysiology or convergence among several complementary evidence channels. The false negative cases revealed that PRIORITI failed in cases where a mismatch between a single-target, pair-level representation and an asset whose efficacy depends on multi-target pharmacology, an intermediate phenotype and a selected patient population. More generally, clinical success can emerge through mechanisms that are not fully contained within a T-I pair, including pleiotropic physiology, modality-specific effects, patient stratification and indication reframing.

Our study also has several limitations and highlights open questions for future work. First, although the LLM protocol explicitly excludes drug-, trial- and regulatory-level information during evidence synthesis, the complete absence of leakage cannot be guaranteed in principle for any model that may contain latent knowledge of prior development outcomes. ^40^ We therefore conducted post hoc checks. Systematic review of the LLM-synthesized evidence did not reveal instances of drug- or trial-level leakage (**Supplementary Note 2**), supporting the view that simple outcome recall was unlikely to be the dominant source of signal.

Second, the programme-derived outcome is an imperfect proxy for biological validity. A T-I pair was defined as successful if at least one associated programme reached launch-like status, and unsuccessful otherwise. This label is pragmatic and reproducible, but it conflates biological validity with many downstream determinants, including molecule quality, modality, dosing, formulation, therapeutic index, commercial decisions, sponsor strategy, endpoint selection and trial execution. A biologically sound T-I hypothesis can fail because the wrong asset, dose, patient population or endpoint was chosen; conversely, a weak direct T-I rationale can succeed because the drug acts through broader physiology or an intermediate phenotype. These considerations imply that PRIORITI should be interpreted as estimating translational biological support, not the causal probability that target modulation alone will produce regulatory success.

Third, the current framework is intentionally non-agentic. It performs constrained evidence synthesis followed by supervised prediction, but does not yet exploit self-evolving AI workflows with long-term memory^41^, self-critique^42^ or adaptive error correction^43^. Such systems could, in principle, improve the handling of conflicting evidence, edge cases and recurrent failure modes. Whether these agentic capabilities can be incorporated without compromising auditability, calibration or leakage control remains an open question.

We conclude the study with two potential complementary paths. The first is live prospective validation through a continuously updated, prediction-locked registry of T-I hypotheses containing public time stamps, frozen evidence packets, prespecified model scores and subsequent outcome tracking. The second is richer, source-grounded reasoning evidence synthesis using time-indexed retrieval, claim-level provenance and explicit representation of conflicting evidence.

## METHODS

Methods are available in the Methods section.

Note: One supplementary table file (Supplementary_tables.xlsx), one supplementary figures file (Supplementary_figures.pdf), and one supplementary notes file (Supplemenytary_notes.docx) are available.

## ACKNOWLEDGEMENTS

Research reported in this work was supported by National Cancer Institute under Award Numbers R01CA263494, U01CA293883, and P30CA016672, National Institute of Mental Health under Award Number R01MH136055; National Institute on Aging under Award Numbers RF1AG082938 and R01AG085581; and The U Foundation U-Pilot Award. The content is solely the responsibility of the authors and does not necessarily represent the official views of the National Institutes of Health.

## AUTHOR CONTRIBUTIONS

C.W., B.Z., and W.Z., designed the study. W.Z. and C.W. developed the PRIORITI model. W.Z. curated and analyzed the data. Z.Z., L.W., M.L. helped with preparing data resources, data analysis, and provide feedback on study and platform design. W.Z., C.W., and B.Z. wrote the manuscript with feedback from all authors.

## CLINICAL TRIAL NUMBER

Not applicable.

## Methods

### Data Acquisition and Cohort Definition

Citeline Pharmaprojects records were extracted as a single database snapshot on December 5, 2025. We programmatically parsed drug-level records to extract specific target genes, mapping Target Entrez Gene IDs to official gene symbols, and standardized disease indications by mapping Citeline disease terms to canonical National Institutes of Health Medical Subject Headings (MeSH)^44^. We rigorously filtered out combination therapies and diagnostic indications. Furthermore, we excluded programs whose highest recorded status was “Preclinical”, retaining only those that had advanced to at least Phase I clinical evaluation. The full derivation of the raw and disease-level cohorts is summarized in **Supplementary Table 1**.

Drug-level development records were subsequently aggregated into unique target-indication (T-I) pairs. For each T-I pair, we determined the maximum clinical development phase achieved across all associated pharmacological assets. We defined the primary outcome label by assigning clinical “Success” to T-I pairs that successfully reached “Launched”, “Registered”, or “Pre-registration” status. Conversely, we classified programs concluding development at Phase I, Phase II, or Phase III, as well as those explicitly marked as suspended, discontinued, or withdrawn, as “Failures”. To ensure data fidelity, ambiguous historical events occurring prior to the year 2000 were systematically excluded.

We then retrospectively reconstructed temporally separated cohorts using 1 October 2024 as the index date. The historical cohort retained launch-like pairs only if launch year was before 2025 and retained all other pairs only if event date was before October 1, 2024. This pre-cutoff pool was then restricted to T-I pairs overlapping the foundational dataset of Minikel et al^8^, providing a mature pre-cutoff benchmark aligned with prior genetics-based forecasting studies. The resulting historical cohort was split into training and held-out test sets using a random 80/20 partition with seed 42. Stepwise derivation and the train-test split are summarized in **Supplementary Table 2.**

To evaluate genuine out-of-time forecasting, we then constructed a post-October-2024 benchmark from Citeline T-I pairs whose outcome was resolved after the LLM knowledge cutoff (**Supplementary Table 3**). At the aggregated pair level, a post-cutoff positive was defined as a pair reaching “Launched”, “Registered”, or “Pre-registration” status with “Launch Year” ≥ 2025, whereas a post-cutoff negative was defined as a non-launch pair with a qualifying update date on or after October 1, 2024. We first removed locally invalid or irrelevant entries, then retained only inactive programs or launch-like programs and finally removed the pairs already present in Minikel^8^ to obtain the new-only benchmark pool. We then manually curated this set to remove pairs that did not represent clean prospective cases relative to the cutoff (**Supplementary Note 1**). Human curation was necessary because some records in the database lacked informative Launch Year annotations, and some pairs required inspection of detailed event histories to determine whether the apparent post-cutoff outcome actually reflected pre-cutoff approval- or launch-related activity, or instead represented a later administrative update to an already established program history. Specifically, we excluded 29 pairs with final positive labels for which matched raw Citeline history already contained pre-cutoff success evidence, indicating that the pair had launched before the cutoff. We also excluded 26 final-negative pairs for which manual review showed that development had already clearly ceased before the cutoff and the post-cutoff event did not represent renewed development activity, trial reactivation, or any other label-changing update. The final out-of-time benchmark comprised 765 T-I pairs, including 89 successes and 676 failures.

### LLM Evidence Synthesis

To eliminate outcome circularity and prevent data leakage, we followed a domain-grounded instruction framework and formulated genetics-based evidence extraction as a constrained inference task utilizing a LLM (GPT-5, version GPT-5-2025-08-07). To ensure the model remained strictly blind to recent clinical outcomes, we relied exclusively on its internal knowledge cutoff and explicitly disabled external tool use, such as web search and function calling, forcing generation to rely solely on the model’s internal knowledge. Furthermore, the system prompt explicitly prohibited the model from consulting, using, or citing drug- or clinical trial-related information, including drug names, trial outcomes, regulatory labels, and registries. Full prompt templates are provided in **Supplementary Table 8**.

The model summarized evidence across four prespecified, drug-agnostic channels, each capturing a distinct facet of target–disease support. Human genetic causality (HGC) reflects the strength and directness of human genetic evidence linking the target gene to the indication, drawing on genome-wide and fine-mapped association signals, variant-to-gene links, molecular quantitative-trait-locus (QTL) colocalization, rare-variant burden, and phenome-wide association studies. Biological coherence (BIO_COH) reflects functional-genomics support that the target is mechanistically embedded in disease-relevant biology, including tissue- and cell-type expression, regulatory and epigenomic context, perturbation evidence, pathways, and protein–protein interaction networks. Functional and animal support (FUNC_ANIMAL) reflects experimental and model-organism evidence, such as knockout or knock-in phenotypes evaluated under conservative orthology and phenotype-alignment criteria, that aligns target modulation with the disease phenotype. Cross-source consistency (CONSISTENCY) reflects the degree to which these independent lines of evidence agree, after deduplication and resource-quality weighting to avoid inflated support from mirrored databases.

Each channel was scored on a 0 to 6 scale using a fixed qualitative rubric held invariant across all T-I pairs, with higher scores denoting stronger, more direct, and better-corroborated evidence. The four channel scores were then integrated into an overall integrated causal evidence score (ICES; 0 to 6), which was mapped to five prespecified verdict levels (insufficient, weak, moderate, strong, and very strong) to enable consistent calibration across targets and indications; following the source framework, ICES ≥ 4.0 denoted strong or very strong integrated causal evidence. To ensure transparency and auditability, the protocol enforced an evidence-ledger (“cite or omit”) principle, in which every substantive causal or direction-of-effect claim was linked to at least one indexed primary study or curated database record, and contradictory evidence was contextualized rather than discarded unless explicitly refuted. All outputs were returned as a single schema-valid JSON record containing the five scalar scores, the channel-specific evidence summaries, an evidence register of supporting sources, and an integrated rationale.

### Feature construction

Each T-I pair was represented using four complementary feature blocks designed to capture both structured evidence strength and richer semantic biological context. First, we used the five scalar LLM-derived evidence scores, comprising the four channel-level evidence scores, including human genetic causality (HGC), biological coherence (BIO_COH), functional and animal support (FUNC_ANIMAL), and cross-source consistency (CONSISTENCY), along with the integrated causal evidence score (ICES). These variables provided a compact quantitative summary of the LLM-synthesized evidence profile.

Second, to preserve information not captured by scalar scores alone, we constructed LLM-derived embeddings from the model-generated natural-language rationales. For each T-I pair, the rationales corresponding to the evidence channels were concatenated into a single text representation and embedded using gemini-embedding-001, yielding a 768-dimensional LLM-derived embedding. This representation was designed to retain higher-order semantic structure in the synthesized biological argument, including how supporting, conflicting, and uncertain evidence was organized across channels.

Third, to encode target identity independently of the LLM reasoning step, we generated a 512-dimensional gene summary embedding from functional gene summaries retrieved from MyGene.info^45^. Fourth, to encode disease identity in a parallel manner, we generated a 512-dimensional indication definition embedding using indication definitions and scope notes obtained from Medical Subject Headings (MeSH)^44^. The dimensionalities of the static entity-context blocks were chosen to satisfy the input-size constraints (< 2000 features) of the AutoGluon v1.5.0 and configuration used in this study.

The final feature representation for each T-I pair therefore consisted of 1,797 total features, obtained by concatenating the 5 scalar LLM-derived evidence scores, the 768-dimensional LLM-derived embedding, the 512-dimensional gene summary embedding, and the 512-dimensional indication definition embedding. This four-block design allowed the downstream supervised model to integrate compact quantitative evidence summaries with richer semantic representations of the synthesized biological rationale, target biology, and indication context.

### Predictive modelling and evaluation

Using the feature representation described above, we formulated T-I outcome forecasting as a supervised binary classification task. Model development was conducted exclusively on the historical training set. We used AutoGluon v1.5^23^ to train and compare a diverse set of tabular learning algorithms, including LightGBM^24^, CatBoost^25^, RealMLP^26^, TabDPT^27^ and RealTabPFN-v2.5^28,29^. Because the task is class-imbalanced and the main practical objective is to rank true successful programmes highly, model selection was guided primarily by average precision / area under the precision-recall curve. Training used an intensive bagged cross-validation scheme with 10 folds, allowing each candidate learner to be fitted repeatedly across resampled subsets of the training data to improve stability and reduce overfitting. AutoGluon then combined the resulting base learners into a weighted ensemble, in which the final predicted probability was obtained as a learned combination of the individual model outputs. Unless otherwise stated, this weighted ensemble was used as the primary model throughout the study.

After training, the final locked model was evaluated without refitting on two downstream datasets: (i) a strictly held-out historical test set for in-distribution evaluation and (ii) an independent post-cutoff future benchmark for out-of-time evaluation under temporal separation. Model performance was summarized using both threshold-free and threshold-dependent metrics. Threshold-free assessment included ROC-AUC, PR-AUC, and expected calibration error (ECE), which together capture discrimination and probability calibration. Because the practical goal of PRIORITI is to prioritize a small subset of high-probability T-I hypotheses for downstream evaluation, we additionally reported precision among the top-ranked predictions by model probability, including Precision@3%. Threshold-dependent metrics included accuracy, balanced accuracy, precision, recall, F1-score, and Matthews correlation coefficient (MCC)^46^. We selected a single operating threshold using training data alone by maximizing F1-score on cross-validated training predictions, and then fixed this threshold for all downstream evaluations.

To assess whether LLM-derived biological rationales added value beyond static target and indication context, we constructed a separate static entity-context baseline. This baseline excluded all LLM-derived evidence scores, rationales and evidence summaries, and represented each T-I pair using only gene-summary embeddings from MyGene.info and indication-definition embeddings from MeSH disease definitions and scope notes. The static baseline used 512-dimensional embeddings for both entity types, which are consistent with the dimensions used in PRIORITI. The resulting baseline contained 1,024 features per T-I pair and was trained and evaluated with the same supervised modelling and evaluation protocol as the full model.

### Entity-disjoint evaluations

To test whether PRIORITI captured transferable target-disease relationships rather than recurrent entity identities, we constructed three progressively stricter identity-disjoint re-partitions of the same historical cohort, in each case removing exact entity overlap between training and test.

For the target-disjoint split, the set of unique target genes was randomly partitioned into disjoint training and held-out groups (NumPy RandomState seed 42), and each T-I pair was assigned according to its target, so that no target gene in the test set appeared during training. This yielded 6,917 training and 1,563 test pairs, with a test-set success prevalence of 19.0%. The indication-disjoint split was constructed analogously by partitioning unique indications so that no test indication was observed during training, yielding 6,270 training and 2,210 test pairs (prevalence 13.8%). For the doubly-disjoint split, target genes and MeSH indications were partitioned independently, assigning approximately 30% of genes and 30% of indications to the held-out side (seed 42); only pairs whose target and indication both fell on the training side were retained for training, and only pairs whose target and indication both fell on the held-out side were retained for testing, whereas the two cross quadrants, in which one entity was in training and the other was held out, were discarded. This produced a test set in which neither the target nor the indication of any pair appeared in training, comprising 4,269 training and 678 test pairs (prevalence 11.5%).

Within each split, the full PRIORITI model and the static target and indication context baseline were retrained from scratch on that split’s training pairs and evaluated on its disjoint test pairs, using the identical feature construction, embedding dimensionality and AutoGluon configuration as the primary held-out analysis. Discrimination was summarized by ROC-AUC and PR-AUC with 95% bootstrap confidence intervals (2,000 resamples, seed 42). For the doubly-disjoint split, the difference between the full model and the baseline was additionally assessed by a paired bootstrap over the common test pairs (2,000 resamples), reporting the mean difference in ROC-AUC and PR-AUC with its 95% confidence interval.

### Internal temporal controls

To distinguish temporal distribution shift from potential label leakage as the source of the reduced out-of-time performance, we performed two internal controls conducted entirely within the model’s knowledge horizon, with all programmes resolved on or before the October 2024 cutoff, using the identical prompting, feature-construction and supervised-modelling pipeline as the main model. In the forward control, we trained on programmes whose outcomes resolved before 2022 and evaluated on two sets: a contemporaneous held-out set from the same pre-2022 period, and a forward set of programmes that resolved between 2022 and the October 2024 cutoff. The pre-2022 cohort was partitioned 80:20 into training (N = 5,035) and held-out (N = 1,259; prevalence 0.107) sets, and the forward set comprised 1,912 programmes (prevalence 0.184). In the reverse control, we inverted the eras, training on recent programmes (resolved 2015 to October 2024) and evaluating on earlier programmes (resolved 2000–2014; N = 3,369; prevalence 0.103), whose outcomes are long settled and fully within the model’s knowledge; to provide a matched same-era reference, the 4,837 recent programmes were partitioned 80:20, stratified by outcome, into training (N = 3,869) and held-out (N = 968; prevalence 0.140) sets. Because both evaluation windows in each control precede the knowledge cutoff, neither can benefit from post-cutoff outcome information. ROC-AUC and PR-AUC were estimated on each evaluation set, with 95% confidence intervals from 2,000 bootstrap resamples.

### Ablation analyses

To assess the contribution of each feature block, we performed two complementary analyses. First, we conducted feature-block ablation by training separate AutoGluon models using individual feature blocks or selected combinations of feature blocks. The feature blocks were: five scalar LLM-derived evidence scores, 768-dimensional LLM rationale embeddings, 512-dimensional gene-summary embeddings and 512-dimensional indication-definition embeddings. Each ablation model was trained using the same historical training set and evaluated on the same held-out test set as the full model. Performance was compared using ROC-AUC, PR-AUC, ECE, F1-score, balanced accuracy, and MCC.

Second, we performed grouped permutation importance analysis on the trained full model. Features were grouped into the same four blocks: scalar LLM-derived evidence scores, LLM rationale embeddings, gene-summary embeddings and indication-definition embeddings. For each group, all features within that block were jointly permuted across samples while the remaining feature blocks were left unchanged. We then recalculated PR-AUC on the held-out test set. The decrease in PR-AUC relative to the intact full model was used as the group-level importance score. This analysis was intended to estimate the marginal predictive contribution of each feature block within the fitted full model.

### Sensitivity and robustness

To assess whether PRIORITI performance depended on a specific modelling algorithm, we compared the primary weighted AutoGluon ensemble with the major individual learner families included in the training framework, including LightGBM, RealMLP, TabDPT and RealTabPFN-v2.5. All models used the same feature representation and historical training data and were evaluated on the same held-out historical test set. Performance was summarized using ROC-AUC, PR-AUC and ECE, with 95% confidence intervals estimated by bootstrap resampling.

To evaluate robustness across disease contexts, we stratified the held-out historical test set by therapeutic area. Within each therapeutic-area stratum, we computed ROC-AUC, PR-AUC and outcome prevalence. Because PR-AUC depends strongly on baseline prevalence, we additionally reported fold enrichment, defined as PR-AUC divided by the success prevalence within the same therapeutic-area stratum. Confidence intervals for stratum-specific metrics were estimated using 1,000 bootstrap resamples.

To assess robustness to the choice of text-embedding model, we regenerated all text representations using two alternative embedding backbones and retrained the full model without any other change. The LLM-rationale, gene-summary and indication-definition texts were re-embedded with OpenAI text-embedding-3-large and with the open-weight Qwen3-Embedding-8B (accessed via OpenRouter), in each case matching the dimensionality of the primary model (768 dimensions for the rationale embedding and 512 dimensions for the gene and indication embeddings) and applying the same L2 normalization. Holding the feature set, embedding dimensionality and downstream AutoGluon configuration fixed, each variant was trained on the same historical training set and evaluated on the same held-out historical test set. Performance was summarized using ROC-AUC and PR-AUC with 95% bootstrap confidence intervals.

### Prompt-based benchmark evaluation

We used three direct prompting settings to characterize the role of the LLM on the out-of-time benchmark: a knowledge-recognition setting, a biology-only zero-shot forecasting setting, and an evidence-informed zero-shot forecasting setting. Full prompt templates are shown in **Supplementary Tables 9-11**.

In the knowledge-recognition setting, the LLM received only the target symbol and indication name and was asked to determine whether the pair represented a known success, a known failure, or an unknown outcome. The prompt imposed a high-confidence recognition standard and explicitly instructed the model to default to unknown if it could not confidently recall a specific clinical or regulatory outcome involving direct modulation of the target in the given indication. This setting was used to assess whether the model appeared to claim prior knowledge of benchmark outcomes.

In the biology-only zero-shot forecasting setting, the LLM was asked to predict success or failure for the same T-I pair using only its internal biological and translational knowledge. No LLM-derived evidence summaries or scalar scores were provided. This setting was intended to measure the model’s direct zero-shot forecasting ability in the absence of structured external evidence.

In the evidence-informed zero-shot forecasting setting, the LLM was additionally provided with the five LLM-derived evidence scores and their matched summaries for HGC, BIO_COH, FUNC_ANIMAL, CONSISTENCY, and ICES, and was asked to make the same binary success-versus-failure prediction. This setting was designed to evaluate whether direct LLM forecasting improved when the model was supplied with the same curated translational evidence used by the supervised predictor.

Predictions from the two forecasting settings were evaluated against the out-of-time benchmark labels and compared with the supervised model. The evidence-informed forecasts were also incorporated into a downstream hybrid analysis to assess whether they contributed complementary signal in borderline cases.

### LLM-based case explainer

To support case-level interpretation of PRIORITI predictions, we developed a constrained, label-blinded LLM-based explainer that converted each locked model output into a standardized translational rationale. The explainer was applied only after model training, threshold selection and out-of-time prediction were complete, and was not used to train, calibrate, threshold or modify the predicted probabilities. Its purpose was to make the locked model output interpretable at the T-I level by summarizing why a given pair was prioritized or deprioritized based on the evidence already available to the forecasting model.

For each out-of-time T-I pair, we constructed an explainer packet containing only prespecified information: the target, indication, target description, indication description, PRIORITI predicted probability, cohort-relative rank and percentile, the five LLM-derived evidence scores, and the corresponding channel-level reasoning summaries. The five evidence channels were human genetic causality, biological coherence, functional and animal support, cross-source consistency, and the integrated causal evidence score. The packet did not include the benchmark label, resolved clinical outcome, trial history, regulatory status, drug-specific information, external literature retrieved after prediction, or any post-cutoff outcome information.

The explainer used a fixed prompt workflow; the full prompt is provided in **Supplementary Table 12**. External tools, web search and iterative retrieval were disabled. The prompt instructed GPT-5 to rely only on the supplied packet and to generate a concise, decision-oriented rationale describing how the locked model prediction should be interpreted for T-I prioritization. The prompt also instructed the explainer to preserve uncertainty when the evidence was incomplete, indirect or discordant, and not to introduce trial-, regulatory- or drug-specific claims absent from the input packet. The output was constrained to a predefined structured schema including a model prediction summary, enhanced translational rationale, model-added value, key evidence drivers, drug discovery implication, remaining uncertainties and a one-sentence decision takeaway. Automated validation was used to check that outputs matched the requested T-I pair, preserved the supplied evidence scores, and did not contain prohibited outcome, regulatory or drug-specific claims.

We applied this explainer uniformly to all 765 out-of-time T-I pairs. These explainer outputs were used only for case-level interpretation and auditability, not for quantitative model evaluation. This design allowed the case-level analysis to assess whether PRIORITI could provide auditable, decision-facing biological rationales without allowing the explainer input to contain the observed outcome.

## Code availability

The code to reproduce results, along with associated datasets and a basic README for guidance, is provided as Supplementary Material for review; the final version of the code will be archived on GitHub upon publication.

## Data availability

The human expert-curated genetic evidence data can be obtained from https://github.com/ericminikel/genetic_support/. All data generated by the present study can be found on OSF.IO upon publication, which is provided as Supplementary Material for review.

