## Supplementary Figure 1 for "Biological rationales from language models enable leakage-resistant forecasts of target-indication success"

**This PDF file includes:**

Supplementary Figures S1-S5

Supplementary Notes 1-2

**Other Supplementary Materials for this manuscript include the following:**

Supplementary Table S1 (.xlsx) (available in an Excel file)

Supplementary Files with readme (replication codes and data, available in a zip file)

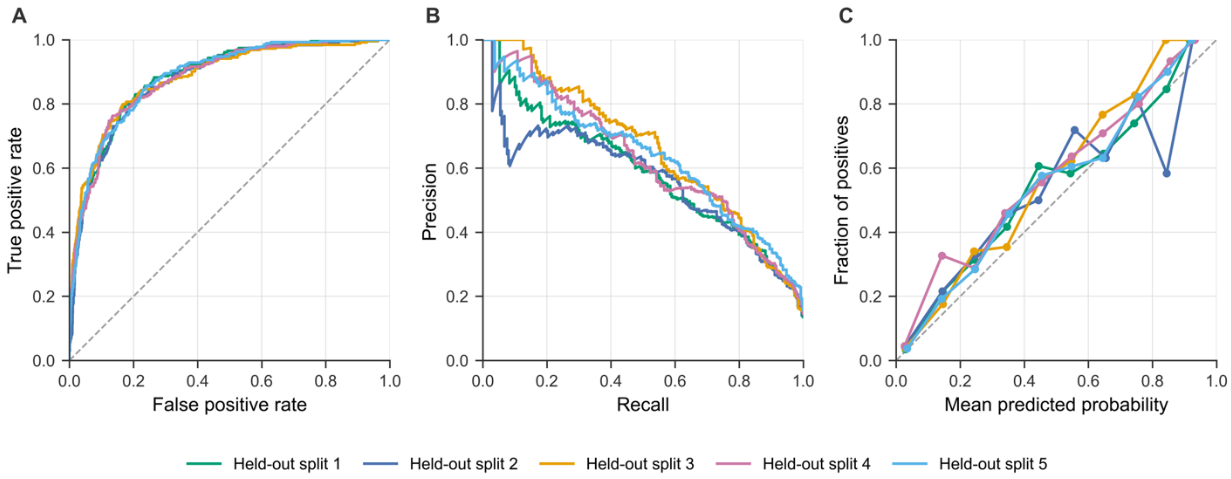


**Supplementary Fig. 1 | Model performance across five independent held-out splits.** **A,** ROC curves for PRIORITI on five independently shuffled held-out test splits. **B,** Corresponding PR curves. **C,** Calibration curves (mean predicted probability vs observed fraction of positives, 10 uniform bins). Held-out split 1 is the original split.

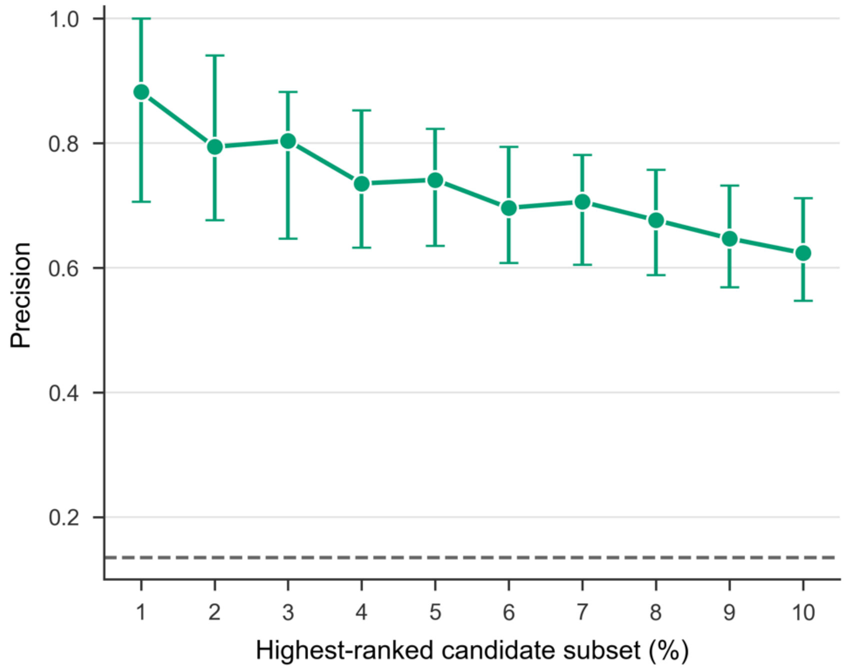


**Supplementary Fig. 2 |** Precision of PRIORITI among the highest-ranked candidate pairs in the historical held-out test set (top 1% to 10%); error bars, 95% bootstrap CIs; dashed line, overall success prevalence.

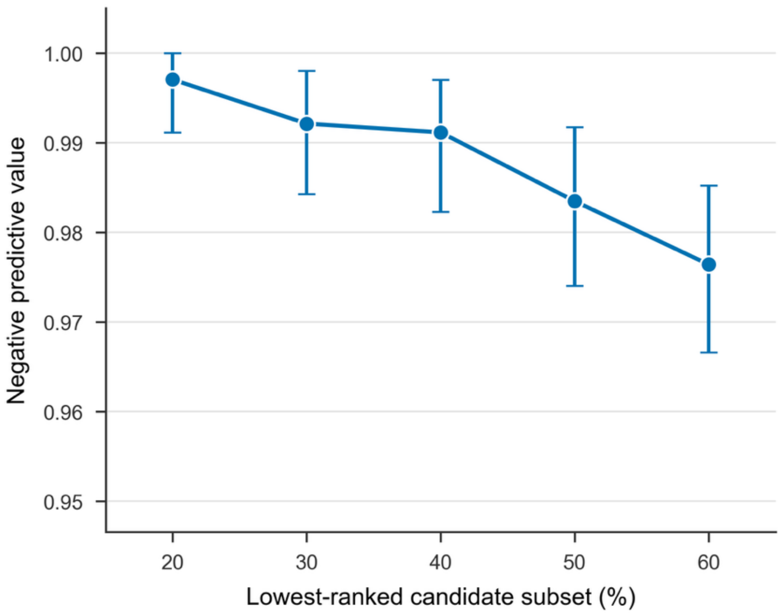


**Supplementary Fig. 3 |** Negative predictive value of PRIORITI among the lowest-ranked candidate pairs (bottom 20% to 60%); error bars, 95% bootstrap CIs; dashed line, overall failure prevalence.

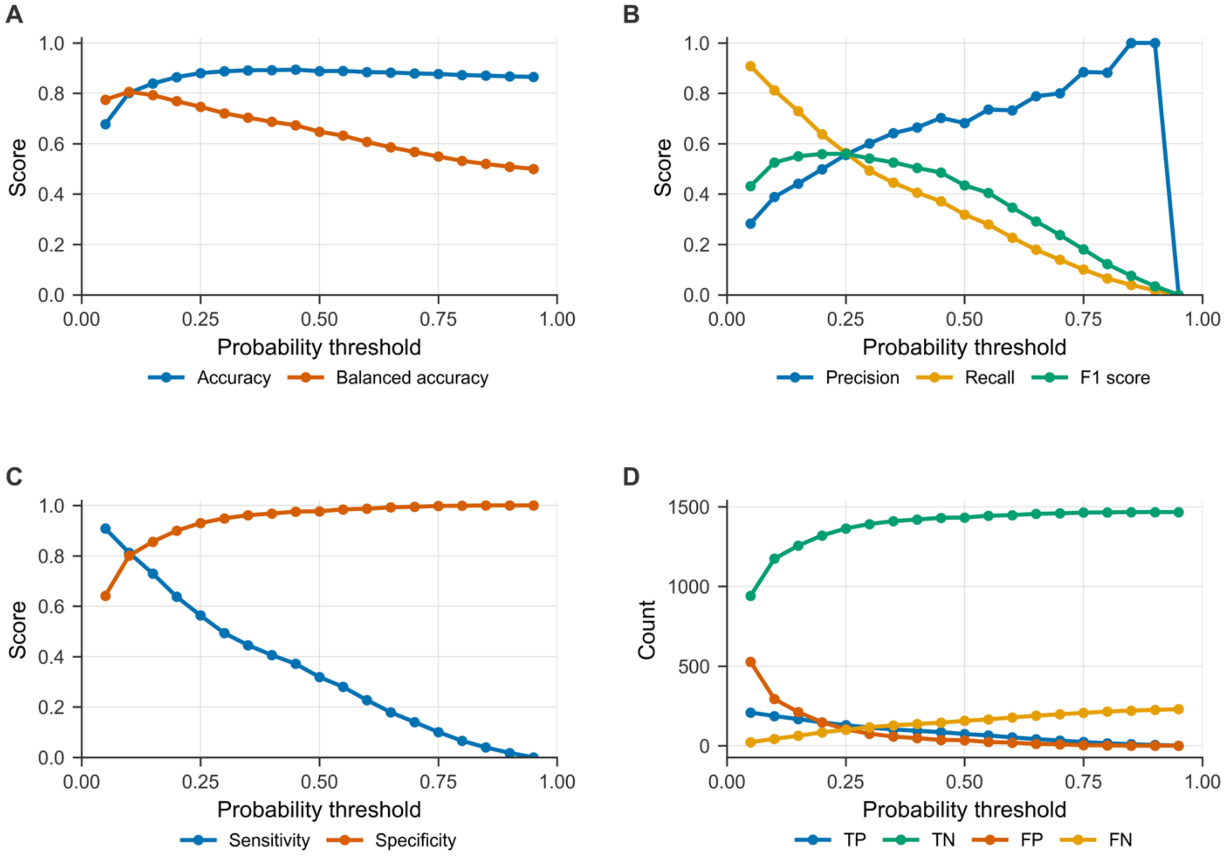


**Supplementary Fig. 4 |** Threshold-dependent classification performance of PRIORITI (thresholds 0.05–0.95). **A,** Accuracy and balanced accuracy. **B,** Precision, recall, F1. **C,** Sensitivity and specificity. **D,** Confusion-matrix components (TP, TN, FP, FN).

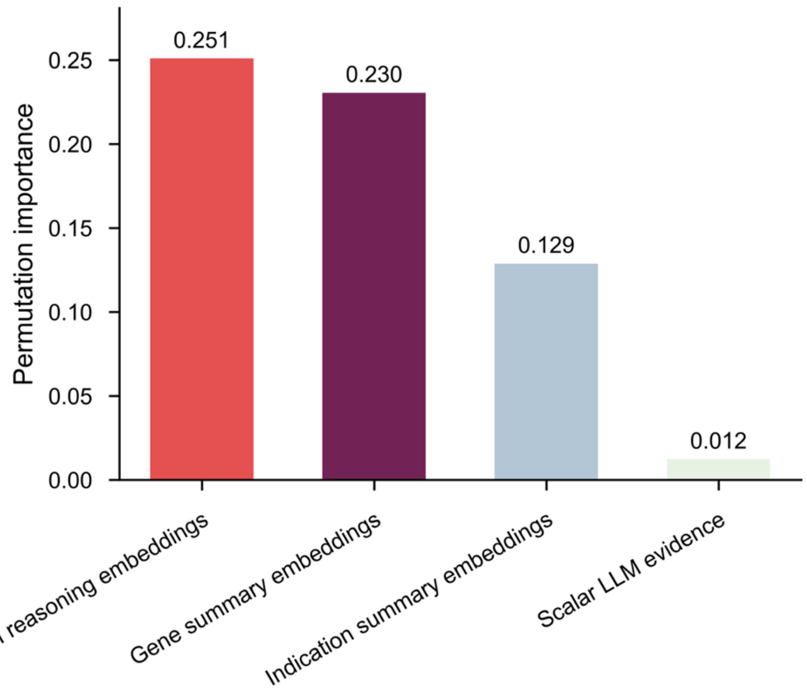


**Supplementary Fig. 5** | Grouped permutation importance of the four feature blocks in PRIORITI on the historical held-out test set (decrease in PR-AUC when each block is permuted): LLM reasoning embeddings largest, then gene- and indication-summary embeddings, with scalar LLM scores contributing little.

### Supplementary Note 1: Post-cutoff benchmark manual curation and leakage control

To ensure that the out-of-time benchmark reflected genuine future-like forecasting rather than latent outcome knowability, we applied additional curation beyond the initial post-October-2024 extraction. Starting from the saved post-2024 metadata snapshot (n = 3,188), we first removed locally invalid or irrelevant entries, then retained only inactive programs or launch-like programs (3,188 to 1,133), and finally restricted the retained set to Minikel indication space (1,133 to 820). The resulting 820-pair benchmark contained 118 positive and 702 negative target-indication pairs. We further performed targeted manual curation to remove pairs whose outcomes were already directionally inferable before the October 2024 cutoff. Human curation was necessary because automated filtering based on summary fields alone was not sufficient to determine whether a pair was truly knowable before the October 2024 cutoff. In particular, some records lacked informative Launch Year annotations, and some pairs required inspection of detailed event histories to determine whether the apparent post-cutoff outcome actually reflected pre-cutoff approval- or launch-related activity, or instead represented a later administrative update to an already established program history.

For this adjudication, GPT-5 was first asked whether each target-indication pair was already known to succeed, already known to fail, or remained unknown. We then compared these claims against matched pre-cutoff Citeline histories. On the positive side, we audited all known_success pairs with ground truth (GT) = 1. Among 39 such pairs, 37 had a success-like final status (Launched, Registered, or Pre-registration), and 29 showed confirmed pre-cutoff success evidence in matched raw Citeline history. Pre-cutoff success evidence was defined as either a disease-level Launch Year ≤ 2024 or a pre-cutoff launch/approval-like event (First Launch, New Launch, First Approval, New Approval, or Supplemental Approval). These 29 pairs were removed from the benchmark. On the negative side, we audited all known_failure pairs with GT = 0. For each pair, we used conservative raw-manual inference from matched disease-level Citeline rows to determine the best-supported pre-cutoff highest non-launch status. A pair was removed only if its final highest status did not exceed this inferred pre-cutoff status and the post-cutoff update was consistent with non-progress or failure. Using this rule, 26 of 57 failure-side candidates were removed, whereas 31 ambiguous cases were retained and no pair was removed on the basis of uncertain evidence. Representative exclusions of target-indication pairs are shown below.

| Category | T-I pair | Final outcome/status | Pre-cutoff evidence | Post-cutoff context | Reason for exclusion |
|---|---|---|---|---|---|

| | | | | | |
|---|---|---|---|---|---|
| Positive-side exclusion | ACLY-D006949 | Registered | Launch Year missing but multiple 2020 approval/launch events already present before cutoff | Still appeared as a success-side pair in the post-cutoff benchmark construction | Already had explicit pre-October-2024 success evidence |
| Positive-side exclusion | ANGPT2-D005128 | Launched | Raw Citeline history already contained First Approval and First Launch events in 2022 | Nominally entered the post-cutoff benchmark because final launch-like resolution was retained there | Success direction was already supported before cutoff |
| Positive-side exclusion | BCL11A-D000755 | Launched | Launch Year 2023 plus multiple 2023-2024 approval events in matched raw history | Final pair remained success-side | Clearly knowable success before cutoff |
| Negative-side exclusion | SSTR1-D018358 | Phase II Clinical Trial | Exact same Gene + MeSH ID pair already existed in the pre-cutoff T-I table at Phase II Clinical Trial | Post-cutoff event was New Approval, but final highest status still remained Phase II Clinical Trial | Exact pair was already present before cutoff and did not advance |
| Negative-side exclusion | MAPK11-D054058 | Phase III Clinical Trial | Raw-manual review supported a pre-cutoff maximum status of Phase III Clinical Trial | Post-cutoff event was Discontinuation Confirmed in 2025; final highest status stayed Phase III Clinical Trial | Post-cutoff update reflected failure/no-progress rather than forward development |
| Negative-side exclusion | ACHE-D057180 | Phase I Clinical Trial | Raw-manual review supported a pre-cutoff maximum status of Phase I Clinical Trial | Post-cutoff event was New Patent in 2025; final highest status stayed Phase I Clinical Trial | No meaningful stage advancement after cutoff |

### Supplementary Note 2: Screening the LLM-derived evidence rationales for clinical-outcome leakage

#### Motivation and definition of leakage

For every target-indication (T-I) pair, a language model (GPT-5, knowledge cutoff October 2024) generates structured genetic and biological evidence across five channels (HGC, BIO_COH, FUNC_ANIMAL, CONSISTENCY, ICES). The free-text summaries of these channels are concatenated, embedded (768 dimensions), and used directly as model features. If a rationale inadvertently revealed the clinical outcome of a pair, the classifier could exploit that disclosure instead of the genetic evidence, inflating apparent performance. The prediction label is defined by that clinical outcome:

**Success:** at least one therapy that directly modulates the target (binding, direct inhibition, or activation) achieved regulatory approval or reached market for the indication.

**Failure:** such a directly-modulating therapy entered clinical development (Phase 1 or later) but was discontinued for lack of efficacy or for safety in the indication.

Outcome leakage therefore means a rationale that reveals or strongly implies that outcome. The generation prompt already forbids the model from consulting, using, or mentioning drugs, therapeutic modalities, or clinical-trial design/results/endpoints/approvals; this note audits whether that instruction held across every pair used in the study.

#### Rationales audited

We audited all 9,245 T-I pairs used in the study across three cohorts, each with five evidence channels, for a total of 46,225 rationale texts. Cohort membership follows the primary split for the historical data and the 765-pair post-cutoff Citeline v4 set for the out-of-time (OOT) cohort.

| Cohort | Pairs | Rationale channels |
|---|---|---|
| Historical train | 6,784 | 33,920 |
| Historical test | 1,696 | 8,480 |
| OOT (Citeline v4) | 765 | 3,825 |
| Total | 9,245 | 46,225 |

#### Audit method

We used a three-layer, reproducible screen that moves from the presence of suspicious wording to whether any such wording could actually predict the label. First, a deterministic lexical screen scans every rationale with a case-insensitive, whole-word regular-expression dictionary of drug- and trial-related terms, capturing a 100-word context window around each match; the dictionary targets outcome disclosure via drug-name morphology (International Nonproprietary Name stems such as -mab, -nib, -vastatin), clinical-trial identifiers (NCT numbers), and regulatory or outcome verbs (approved, launched, withdrawn, discontinued, failed phase, missed or met primary endpoint, topline). Second, an independent language-model judge (GPT-5.5) reads every flagged channel and scores it against the label definition on a four-point severity scale: 0, benign biological terminology; 1, a drug or trial is noted to exist (or is explicitly excluded) but no outcome is revealed; 2, an indirect or ambiguous outcome hint; 3, explicit disclosure that a therapy directly modulating this target for this indication was approved/launched or failed/discontinued. To confirm the lexical screen does not miss paraphrased (keyword-free) leakage, the judge also scored a random sample of unflagged channels as a negative control. Third, because the presence of a keyword is only harmful if it correlates with the label, we test each flag indicator against the label (odds ratio with a 2,000-sample bootstrap 95% confidence interval, seed 42, and Fisher's exact test) and fit a logistic model predicting the label from the flag indicators alone (five-fold cross-validated ROC-AUC); an AUC near 0.5 means the flags carry no label information.

### Results: lexical screen

Only 311 of 46,225 rationale channels (0.7%) contained any drug- or trial-related keyword. The drug-name (INN-stem) detector matched 0 named compounds and there were 0 clinical-trial identifiers (NCT numbers) in the entire corpus. The matches are dominated by generic biomedical wording; the most frequent matched terms are shown below.

| Matched term | Hits | Typical meaning in context |
|---|---|---|
| drug | 109 | P-glycoprotein / xenobiotic transporter annotation |
| mg | 101 | magnesium ion or units, not dosing |
| subcutaneous | 58 | adipose tissue depot, not administration route |
| withdrawal | 49 | opioid/nicotine/ethanol withdrawal phenotype |
| dose | 49 | gene-dosage / dose-response sensitivity |
| interventional | 22 | observational or animal-model study wording |
| overall survival | 18 | observational cohort endpoint |
| therapy | 15 | generic mention, no outcome |
| prospective cohort | 9 | generic biomedical wording |
| phase i | 7 | generic biomedical wording |
| clinical studies | 6 | generic biomedical wording |
| therapeutic | 4 | generic biomedical wording |

### Results: severity adjudication

The judge adjudicated all 311 flagged channels with no errors. The severity distribution was 307 at severity 0 (benign), 4 at severity 1 (existence only), 0 at severity 2, and 0 at severity 3 (explicit outcome disclosure). No channel reached severity 2 or above: the judge found no ambiguous or explicit outcome disclosure anywhere in the corpus.

| Cohort | Flagged | Sev 0 | Sev 1 | Sev 2 | Sev 3 |
|---|---|---|---|---|---|
| Historical train | 205 | 201 | 4 | 0 | 0 |
| Historical test | 76 | 76 | 0 | 0 | 0 |
| OOT (Citeline v4) | 30 | 30 | 0 | 0 | 0 |

By category, 256 channels were benign biological terminology, 51 were non-interventional references (observational or animal studies), and 4 were model self-reflection, in which the model explicitly noted that it excluded interventional or therapy-stratified evidence. The judge additionally identified 16 channels that name a specific drug (L-DOPA, cocaine, levodopa, morphine, naloxone); in every case the compound appears as a disease-inducing exposure or experimental substrate (for example, levodopa within the indication name levodopa-induced dyskinesia, and cocaine, morphine, or naloxone in addiction animal models), never as a therapy modulating the target, and none disclosed a clinical outcome.

As a negative control, the judge scored 300 randomly sampled unflagged channels; 0 reached severity 2 or above and 0 reached severity 3, confirming that the lexical screen does not miss keyword-free leakage.

### Results: do the flags predict the label?

The decisive test is whether any flag is associated with the label. As shown below, the flag indicators are statistically independent of the label in the historical cohorts.

| Flag indicator | Odds ratio [95% CI] | Fisher p |
|---|---|---|
| Any keyword flag | 1.16 [0.75, 1.66] | 0.52 |
| Any outcome-tier flag | 2.35 [0.59, 5.51] | 0.171 |

A logistic model predicting the label from the flag indicators alone achieves ROC-AUC 0.4974 on the historical cohorts and 0.4885 on the OOT cohort, indistinguishable from chance (0.5). The keyword flags therefore carry no exploitable information about the outcome. The 765 OOT pairs resolved after the model's October-2024 knowledge cutoff, so their outcomes could not be known to the model; in this cohort 2.1% of pairs carried any keyword flag and 0 channels reached severity 3.

### Conclusion

Across every rationale used to train and evaluate the model, we find no evidence of exploitable clinical-outcome leakage. Drug- and trial-related wording is rare (under 1% of channels), contains no specific drug names or trial identifiers from the lexical detector, and is overwhelmingly benign biological terminology. An independent language-model judge found no explicit outcome disclosure, and the negative control shows the screen does not miss paraphrased leakage. Most importantly, the presence of any flag is statistically independent of the label and cannot predict it (AUC approximately 0.5), so even the wording that does appear could not have served as a shortcut to the outcome.
